# A functional landscape of chronic kidney disease entities from public transcriptomic data

**DOI:** 10.1101/265447

**Authors:** Ferenc Tajti, Christoph Kuppe, Asier Antoranz, Mahmoud M. Ibrahim, Hyojin Kim, Francesco Ceccarelli, Christian Holland, Hannes Olauson, Jürgen Floege, Leonidas G. Alexopoulos, Rafael Kramann, Julio Saez-Rodriguez

**Affiliations:** RWTH Aachen University, Faculty of Medicine, Joint Research Centre for Computational Biomedicine (JRC-COMBINE), Aachen, Germany; Division of Nephrology and Clinical Immunology, Faculty of Medicine, RWTH Aachen University, Aachen, Germany; National Technical University of Athens, Greece; ProtATonce Ltd, Athens, Greece; Division of Renal Medicine, Department of Clinical Science, Intervention and Technology, Karolinska Institutet, Stockholm, Sweden; Institute for Computational Biomedicine, Heidelberg University, Faculty of Medicine, Bioquant, Heidelberg, Germany

**Keywords:** transcription factor, signaling pathway, CKD, drug repositioning

## Abstract

To develop efficient therapies and identify novel early biomarkers for chronic kidney disease an understanding of the molecular mechanisms orchestrating it is essential. We here set out to understand how differences in CKD origin are reflected in gene expression. To this end, we integrated publicly available human glomerular microarray gene expression data for nine kidney disease entities that account for a majority of CKD worldwide. We included data from five distinct studies and compared glomerular gene expression profiles to that of non-tumor parts of kidney cancer nephrectomy tissues. A major challenge was the integration of the data from different sources, platforms and conditions, that we mitigated with a bespoke stringent procedure. This allowed us to perform a global transcriptome-based delineation of different kidney disease entities, obtaining a landscape of their similarities and differences based on the genes that acquire a consistent differential expression between each kidney disease entity and nephrectomy tissue. Furthermore, we derived functional insights by inferring activity of signaling pathways and transcription factors from the collected gene expression data, and identified potential drug candidates based on expression signature matching. We validated representative findings by immunostaining in human kidney biopsies indicating e.g. that the transcription factor FOXM1 is significantly and specifically expressed in parietal epithelial cells in RPGN whereas not expressed in control kidney tissue. These results provide a foundation to comprehend the specific molecular mechanisms underlying different kidney disease entities, that can pave the way to identify biomarkers and potential therapeutic targets. To facilitate this, we provide our results as a free interactive web application: https://saezlab.shinyapps.io/ckd_landscape/.

**Translational Statement:** Chronic kidney disease is a combination of entities with different etiologies. We integrate and analyse transcriptomics analysis of glomerular from different entities to dissect their different pathophysiology, what might help to identify novel entity-specific therapeutic targets.

## 1. Introduction

Chronic Kidney Disease (CKD) is a major public health burden affecting more than 10 % of the population globally ^1^ There is no specific therapy and the associated costs are enormous ^2^. The origin of CKD is heterogenous and has slowly changed in recent years due to an aging population with increased number of patients with hypertension and diabetes. Major contributors to worldwide CKD include Diabetic nephropathy (DN) and Hypertensive nephropathy (HN). Other contributors are immune diseases such as Lupus Nephritis (LN) and glomerulonephritides including IgA nephropathy (IgAN), Membranous glomerulonephritis (MGN), Minimal Change Disease (MCD) as well as Focal Segmental Glomerulosclerosis (FSGS) and Rapidly progressive glomerulonephritis (RPGN).

Regardless of the type of initial injury to the kidney the stereotypic response to chronic repetitive injury is scar formation with subsequent kidney functional decline. Scars form in the tubulointerstitium as tubulointerstitial fibrosis and in the glomerulus as glomerulosclerosis. Despite this stereotypic response the initiating stimuli are quite heterogeneous, ranging from an auto-immunological process in LN to poorly controlled blood glucose levels in DN. A better understanding of similarities and differences in the complex molecular process orchestrating disease initiation and progression will guide the development of novel targeted therapeutics.

A powerful tool to understand and model the molecular basis of diseases is the analysis of genome-wide gene expression data. This has been applied in the context of various kidney diseases contributing to CKD ^3–7^, and most studies are available in the online resource NephroSeq. However, to the best of our knowledge, no study so far has combined these data sets to build a comprehensive landscape of the molecular alterations underlying different kidney diseases that account for the majority of CKD cases. We collected data from five large studies with microarray gene expression data from kidney biopsies of patients of eight different glomerular disease entities leading to CKD (from hereon referred to as CKD entities), FSGS, MCD, IgAN, LN, MGN, DN, HN and RPGN. We normalized the data with a bespoke stringent procedure, which allowed us to study the similarities and differences among these entities in terms of deregulated genes, pathways, and transcription factors, as well as to identify drugs that revert their expression signatures and thereby might be useful to treat them.

## 2. Results

### 2.1. Assembly of a pan-CKD collection of patient gene expression profiles

We searched in Nephroseq (www.nephroseq.org) and Gene Expression Omnibus (GEO) ^8,9^ and identified five studies - GSE20602 ^10^; GSE32591 ^11^; GSE37460 ^11^; GSE47183 ^12,13^; GSE50469 ^14^ (see section 4.1.) - with human microarray gene expression data for nine different glomerular disease entities: FSGS, MCD, IgAN, LN, MGN, DN, HN and RPGN, as well as healthy tissue and non-tumor part of kidney cancer nephrectomy tissues as controls (Figure 1A and B). In addition, in one dataset, patients were labeled as an overlap of FSGS and MCD (FSGS-MCD) and we left it as such. These studies were generated in two different microarray platforms. To jointly analyze and compare the different CKD entities, we performed a stringent preprocessing and normalization procedure involving quality control, either cyclic loess normalization or YuGene transformation and a batch effect mitigation procedure (see Methods and Supplementary material). At the end we kept 6289 genes from 199 samples in total. From the two potential controls, healthy tissue and nephrectomies, we chose the latter for further analysis as the batch mitigation removed a large number of genes from the healthy tissue samples.

**Figure 1.**
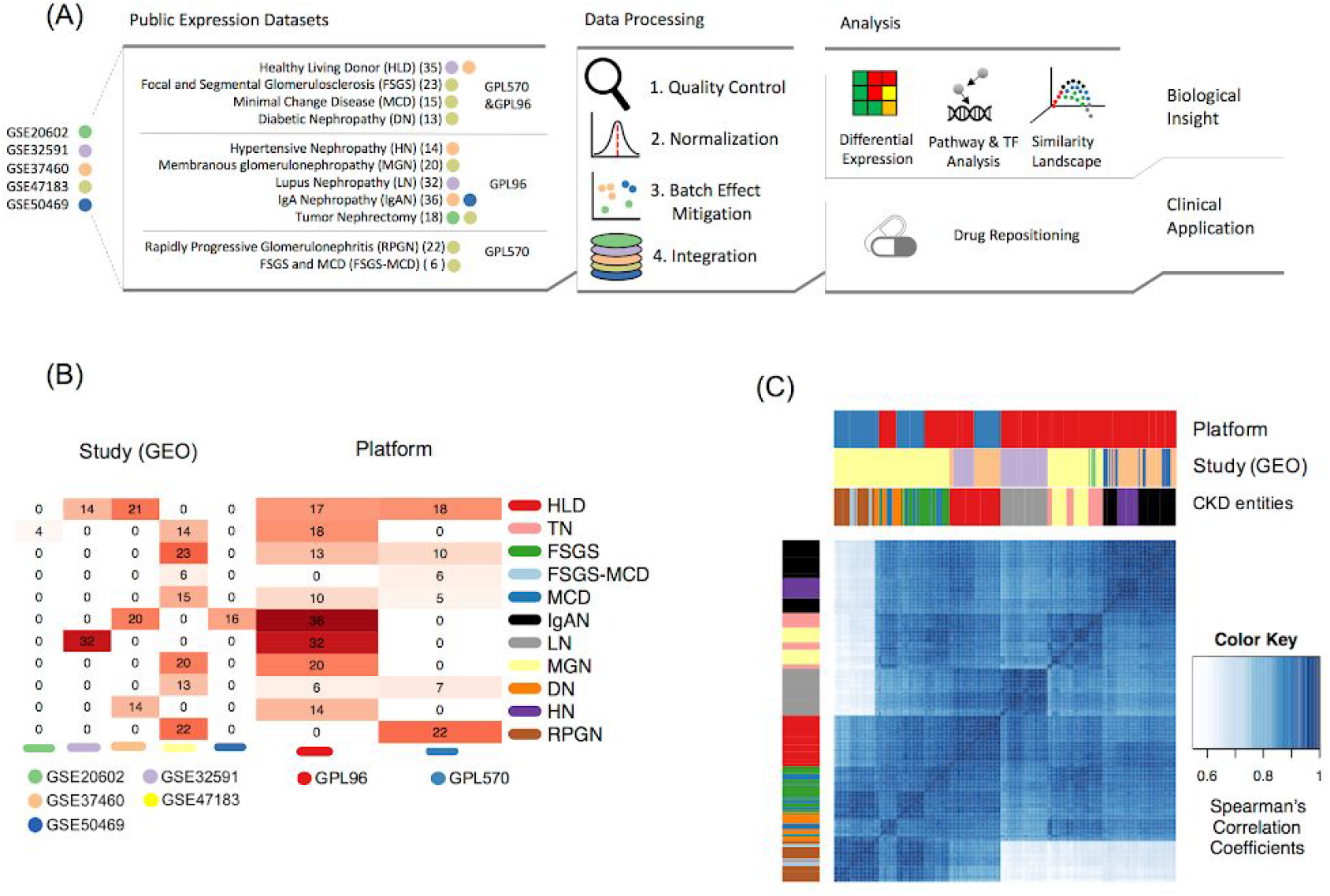
(A) Flow of analysis followed in this study. (B) Heatmap of the distribution of samples across studies and microarray platforms. (C) Hierarchical clustering of the arrays based on gene expression Spearman’s correlation coefficients.

### 2.2. Technical heterogeneity across samples

We first examined the similarities among the samples to assess potential batch effects. Data did not primarily cluster by study source or platform, which can be attributed to our batch mitigation procedure (Figure 1C, Supplementary Figure 1), although some technical sources of variance potentially still remained (see section 4.4. and Supplementary Figure 1). Samples from RPGN and FSGS-MCD conditions seemed to be more affected by platform-specific batch effects than samples from other conditions, due to the unbalanced distribution of samples: RPGN and FSGS-MCD samples were exclusively represented in one study and in one of the two platforms (Affymetrix Human Genome U133 Plus 2.0 Array (GPL570)). Therefore, batch effect mitigation procedure could not be conducted on them.

### 2.3. Biological heterogeneity of CKD entities

We set out to find molecular differences among glomerular CKD entities. First, we calculated the differential expression of individual genes between the different CKD entities and TN using limma ^15,16^. From the 6289 genes included in the integrated dataset, 1791 showed significant differential expression (|logFC| > 1, p-value < 0.05) in at least one CKD entity. RPGN was the CKD entity with the largest number of significantly differentially expressed genes (885), while MCD was the one with the least (75). Twelve genes showed significant differential expression across all the CKD entities (AGMAT, ALB, BHMT2, CALB1, CYP4A11, FOS, HAO2, HMGCS2, MT1F, MT1G, PCK1, SLC6A8). Interestingly, all these genes were underexpressed across all the CKD entities compared to TN. In contrast, QKI and LYZ genes were significantly overexpressed in HN, IgAN, and LN, while significantly underexpressed in FSGS-MCD, and RPGN (and DN for QKI). 107 different genes were significantly differentially expressed relative to TN in at least 6 CKD entities (Figure 2A). Of note, several of the above mentioned genes are considered to be expressed mainly in tubule. This is one drawback of the microdissection technique and future studies using scRNA-seq will dissect which genes are specifically expressed in glomerular cells during homeostasis and disease.

**Figure 2.**
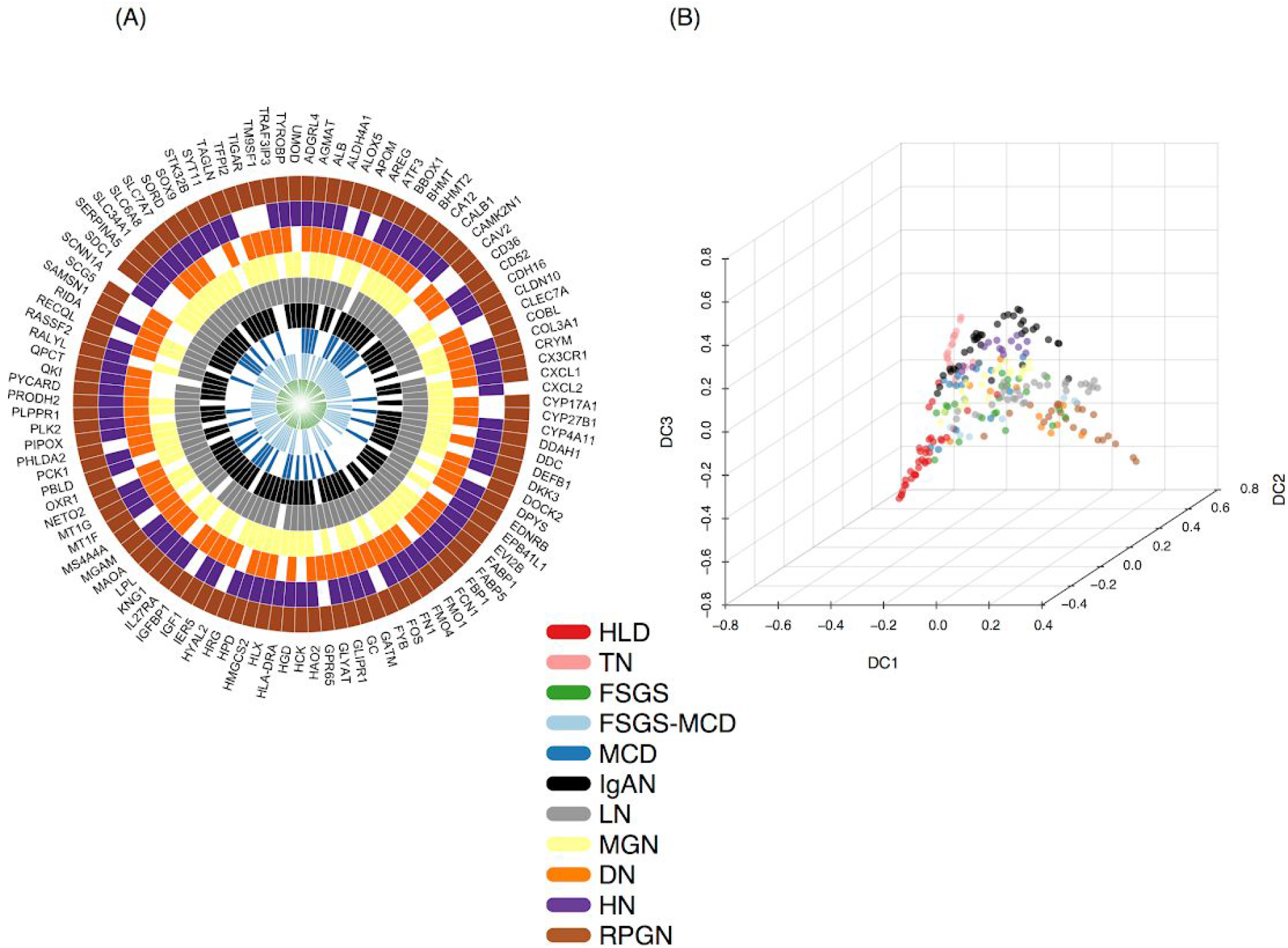
(A) Radial heatmap of consistently differentially expressed genes across six or more disease entities (up- or down-regulation). (B) Diffusion map of CKD entities reveals the underpinning geometric structure of the glomerular CKD transcriptomics data.

To better comprehend the divergence and similarities of the CKD samples, we asked how the distinct CKD entities localised with respect to each other using a common set of differentially expressed genes with regard to the tumor nephrectomies using diffusion maps (Figure 2B). The diffusion distances of each given CKD entity sample relative to tumor nephrectomy samples reflects a non-linear lower dimensional representation of the differences in gene expression profiles between those samples. The Diffusion map orders the patients along a “pseudo-temporal” order, which we interpret here to indicate disease progression severity in glomeruli ^17^.

The most distant condition from nephrectomy samples was RPGN, which is arguably the most drastic kidney disease condition with the most rapid functional decline among the entities included. Interestingly, healthy donor samples were distinct from tumor nephrectomy samples despite the fact that the later were resected distantly from the tumors. This might be explained by either minor contamination with cancer cells or paraneoplastic effects on the non-affected kidney tissue such as immune cell infiltration or solely the fact that the nephrectomy tissue was exposed to short ischemia whereas the biopsy tissue from healthy donors was not. DN and LN were in close proximity to RPGN, whereas HN localised near IgAN. Differences were harder to asses in the middle of the diffusion map, but were visible when plotting the dimension components pair-wise (Sup. Figure 2). For instance, MCD samples spanned from a point proximal to tumor nephrectomy to near FSGS, but some MCD samples were in close proximity to MGN or even hypertensive nephropathy. While it makes sense that MCD as a relatively mild disease with normal light microscopy, is relatively close to the control groups of TN and HLD, it remains unclear why other disease entities such as LN and DN. spread widely in the diffusion map. Unfortunately, the data we used did not include information about disease severity, which might help to explain this heterogeneity with early stage disease possibly closer to the control groups and late stage disease closer to RPGN. Dimension component 1 (DC1) seems to offer a focus on the dissimilarity between the two reference healthy conditions, tumor nephrectomy and healthy living donor from the CKD entities. Dimension component 2 (DC2) provides more insight into the disparity of the reference conditions. Dimension component 3 (DC3) discerns the subtle geometrical manifestation of the distinct CKD entities with regard to each other. In summary, using diffusion maps we find clear differences in the global expression profiles of the CKD entities.

### 2.4. Transcription factor activity in CKD entities

To further characterize the differences among the CKD entities, we performed various functional analyses. First, we assessed the activity of transcription factors (TFs; Figure 3), based the levels of expression of their known putative targets (see Methods). Changes in putative target genes provided superior estimates of the TF activity than the expression level of the transcription factor itself ^18,19^ (Figure 3). We found 10 TFs differentially regulated in at least one CKD entity (Figure 3). Furthermore, we correlated the identified TF’s activities with the expression of those genes, that are encoding for these TFs. The idea was that, while factors with negative correlations are potentially acting as repressors, those with positive correlations are acting as activators. Those with no correlation indicate factors whose activity may be significantly modulated using post-translational modifications or factors whose regulation or expression measurements are unconfident. For instance, Interferon regulatory factor-1 (IRF1) is significantly enriched in LN and moderately correlated-Spearman’s rho (r_s_ = 0.624- with the expression level of the gene encoding for IRF1. This suggests an as of yet undiscovered potential role of IRF1 as a transcriptional activator in LN. In addition, IRF1’s transcriptional activity was elevated in LN compared to the other disease entities. The activity of the upstream stimulatory factor 2 (USF2) - a basic helix-loop-helix (bHLH) TF ^20^ - was estimated to be significantly decreased in MCD compared to the rest of the conditions. Interestingly, USF2’s estimated activity across the CKD entities was inversely correlated - Spearman’s rho (r_s_ = −0.867) - with the expression level of the gene USF2, that is encoding for the TF USF2. Intriguingly, USF2 has been implicated as a potential transcriptional modulator of angiotensin II type 1 receptor (AT1R) - associated protein (ATRAP/Agtrap) in mice ^20^.

**Figure 3.**
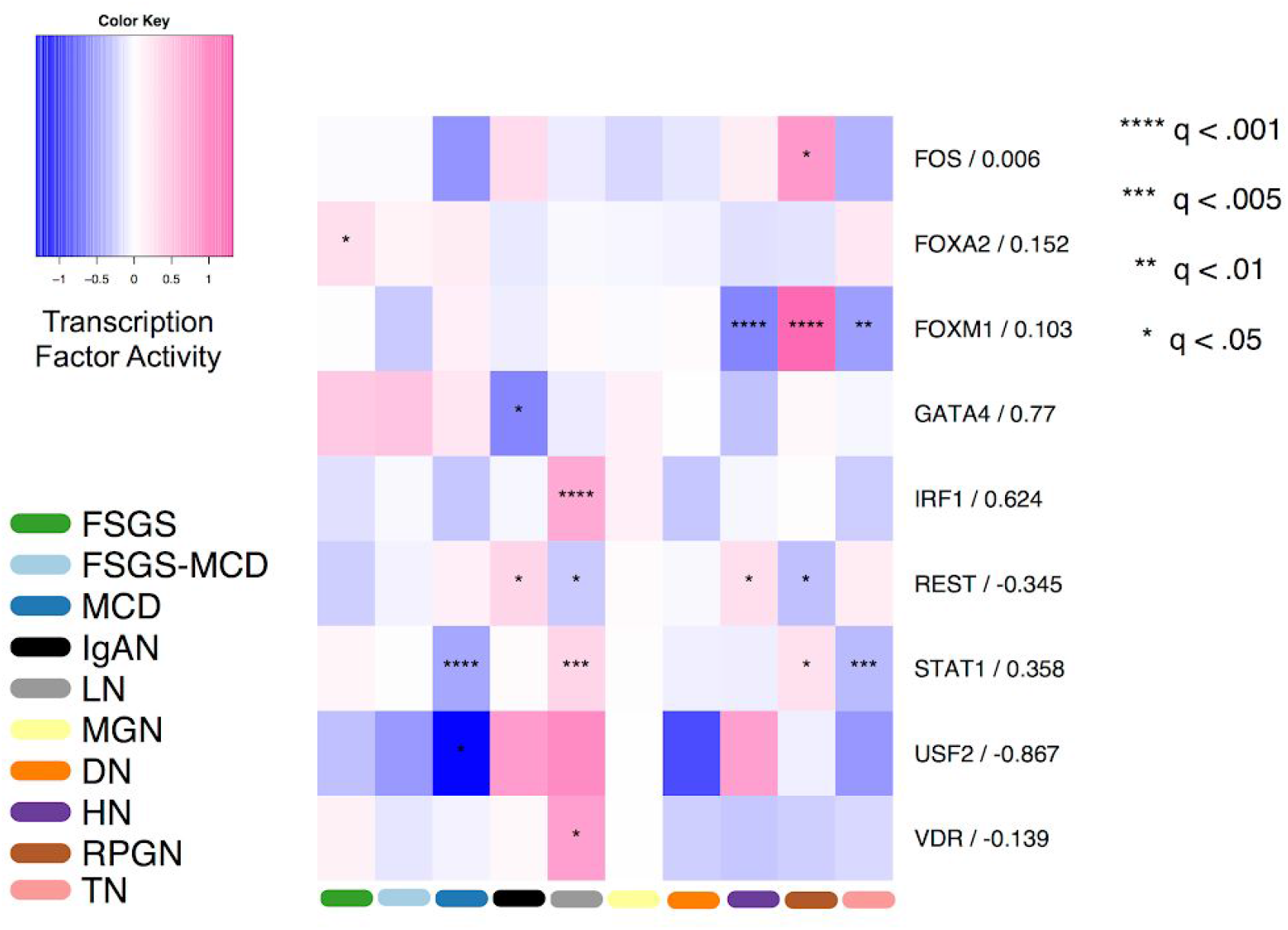
Transcription Factor Activity in Glomerular CKD Entities. Heatmap depicting transcription factor activity (colour) for each CKD entity and tumor nephrectomy in glomerular tissue. Negative numbers (blue) signify decreased transcription factor activity, positive numbers (pink) indicate increased transcription factor activity of an entity relative to the other entities. The corresponding q-value is represented by asterisk(s) (*) to indicate the statistical significance of each TF in each disease entity. The numbers to the right of factor names are Spearman’s rank-based correlation coefficients of factor activity and factor expression across different CKD entities.

We next sought to validate the expression of two identified TFs in human tissue by immunostaining. We stained for USF-2 in human kidney biopsies from healthy controls and patients with MCD. USF-2 was expressed in podocytes, the mainly affected glomerular cell-type in MCD (Figure 4A-B). However, when compared to controls, USF-2 expression in podocytes showed no significant difference detectable by immunofluorescence (Figure 4C-C’’). The reason for this might be that USF-2 activity as a TF might be regulated not only by its abundance in the nucleus but rather by its DNA binding capability in the interaction with other proteins. FOXM1 is a transcription factor of the forkhead box family and a known regulator of cell cycle progression in normal cells as well as a predictor of adverse outcomes across 39 human malignancies ^21^. Our analysis suggests a highly increased activity of FOXM1 in RPGN (Figure 3). We next validated this observation in human biopsy samples from RPGN patients and normal controls. FOXM1 showed a unique expression in CD44 positive glomerular parietal epithelial cells in RPGN lesions whereas we did not find any expression of FOXM1 in healthy human glomeruli (Figure 4. D-F). Consistent with our TF activity analysis, quantification of this finding in 5 RPGN biopsies versus 6 controls yielded a highly significant difference (Figure 4F), indicating that FOXM1 has a significant role in RPGN progression. This data suggest that our computational method might be useful to identify novel regulators in CKD.

**Figure 4.**
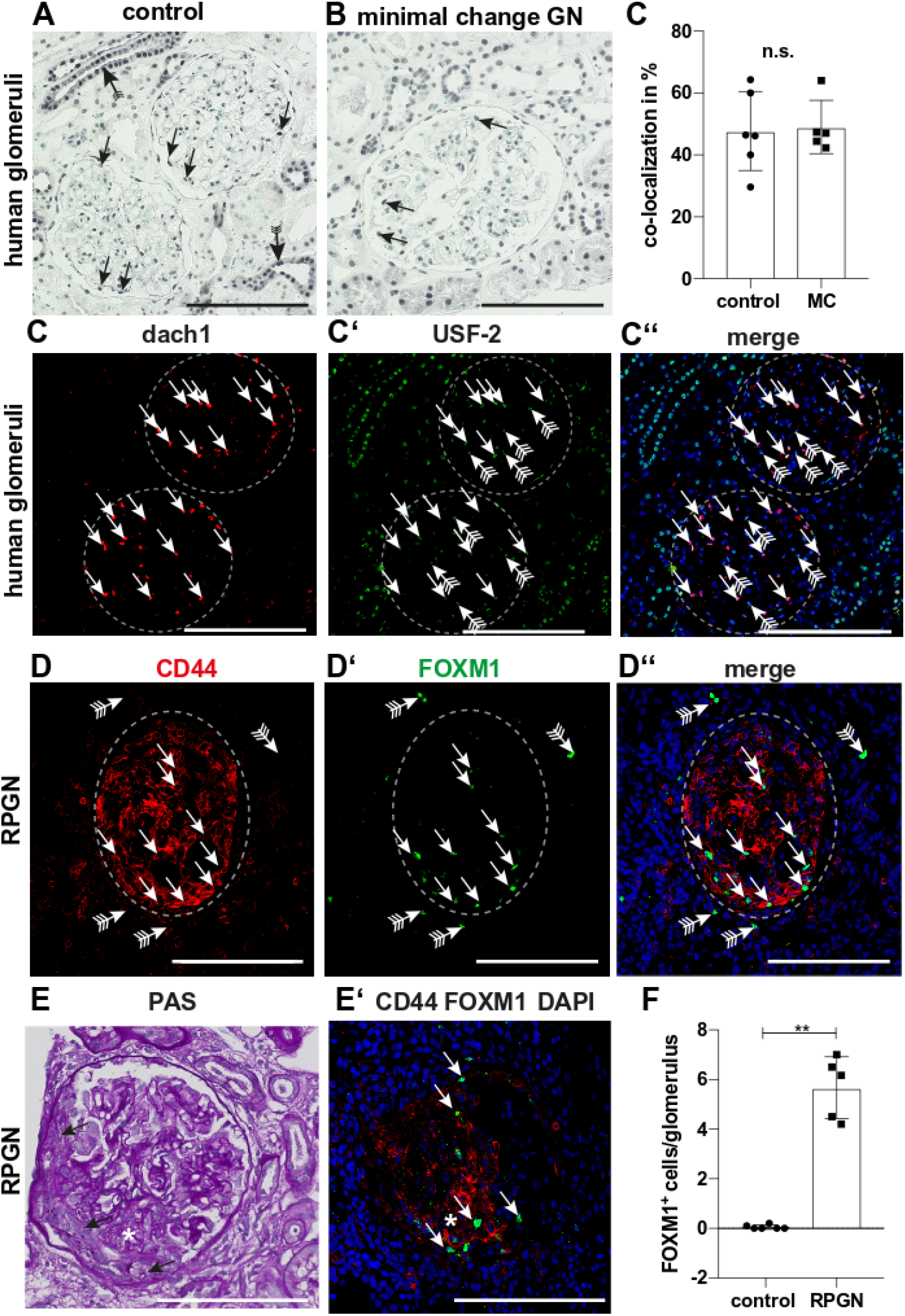
Validation of USF2 and FOXM1 in human kidney biopsies. (**A-C**) Histological validation of USF-2 expression in human biopsies from minimal change disease patients (n=5) and controls (n=6). (**A**) Immunohistochemical staining of USF-2 showed expression in nuclei of many cell types of the kidney included tubular cells (strongest in collecting duct, arrow with tails). In the glomeruli USF-2 expression could be detected in podocytes (arrows). (**B**) USF-2 staining in biopsies from patients with minimal change disease demonstrated a similar staining pattern compared to controls including expression in podocytes (arrows).(**C-C’’**) Quantification of USF-2 expression in podocytes by double-immunofluorescence staining. Co-localized dach1 (podocyte marker in red) and USF-2 (in green). (**D-F**) Histological validation of FoxM1 expression in human biopsies from patients with PRGN (n=5) and controls (n=6). FoxM1 expression was detected most abundantly in glomeruli with crescentic CD44^+^ lesions (arrows in D-D’’). Rarely expression could be noted in the tubular compartment (arrows with tails). (**E-E’**) Serial sections revealed that FoxM1 expression was mainly detected in CD44^+^ cells in the glomerular proliferative lesions (arrows in E). (**F**) Quantification of number of glomerular FoxM1^+^ cells control vs. RPGN (p=0.0043). n.s. not significant, *P < 0.05 by unpaired Mann-Whitney t-test (C and F). Data represents mean ±SD. Scale bars: 100 μm.

### 2.4. Signaling Pathway Analysis

We complemented the functional characterization of transcription factor activities with an estimation of pathways activities with the tools PROGENy ^22^ and Piano ^23^.

#### 2.4.1. Pathway activity of CKD entities using PROGENy

PROGENy infers pathway activity by looking at the changes in levels of the genes affected by perturbation of pathways. This provides a better proxy of pathway activity than assessing the genes in the actual pathway ^22^. We used PROGENy scores to estimate pathway activity in a disease entity from the gene expression data (Figure 5A). Essentially, the degree of pathway deregulation was associated with the degree of disease severity, and present rather divergent activities across the CKD entities. For example, VEGF was estimated to be significantly influential in five CKD entities: RPGN, HN, DN, LN and IgAN, out of which VEGF is predicted to be deactivated in RPGN and DN, but more prominently activated in HN, LN and IgAN. 10 out of 11 pathways were predicted to be significantly deregulated in RPGN with respect to TN, which is aligned with the diffusion map (Figure 2B) outcome; the divergence of RPGN from TN (control) was considerably more prominent both at a global transcriptome landscape and signaling pathway level. Intriguingly, the pathway JAK-STAT did not appear to be affected in RPGN, but was considerably activated in LN and markedly deactivated in DN in comparison to TN. Overall, the separate CKD entities were characterised by distinct combinations, magnitudes and directions of signaling pathway activities according to PROGENy.

**Figure 5.**
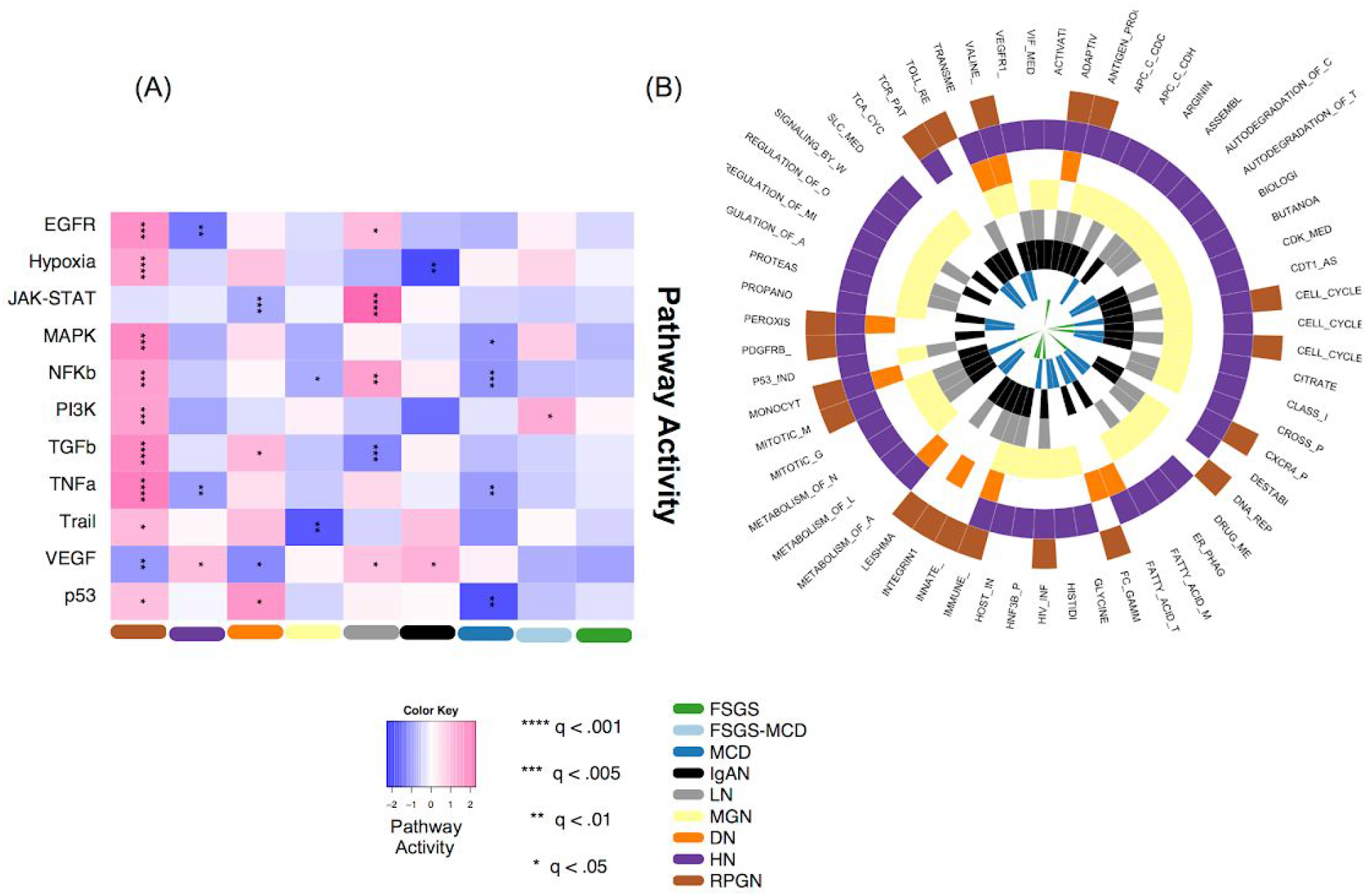
Pathway activity alterations in CKD entities. (A) Heatmap depicting pathway activity (colour) for each CKD entity relative to tumor nephrectomy in glomerular tissue, according to PROGENy ^22^. The magnitude and direction - positive or negative - of PROGENy scores indicates the degree of pathway deregulation in a given CKD entity with regard to the reference condition, tumor nephrectomy. Permutation q-values are used to indicate statistical significance of each pathway in each disease entity, indicated by asterisk(s) (*). (B) Radial heatmap of consensually enriched pathways across three or more disease entities (up-, down-, or non-directional-regulation) according to PIANO ^23^ using MSigDB-C2-CP gene sets.

#### 2.4.2. Pathway enrichment with Piano

While PROGENy can give accurate estimates of pathway activity, it is limited to 11 pathways for which robust signatures could be generated ^22^. To get a more global picture, we complemented that analysis with a gene-set-enrichment analysis using Piano ^23^. A total of 160 pathways out of 1329 were significantly enriched (up-/down-regulated, corrected p-value < 0.05) in at least one CKD entity. HN was the entity with the largest number of differentially enriched pathways (81, 25 down-regulated, 56 up-regulated), while FSGS-MCD did not show significant enrichment for any pathway. Cell-cycle and immune-system related pathways were significantly up-regulated in 7/9 CKD entities (FSGS, HN, IgAN, LN, MGN and RPGN in both cases, DN for immune system, and MCD for cell-cycle); in contrast, the VEGF pathway was differentially enriched in LN only. Interestingly, the TNFR2 pathway was differentially enriched in IgAN, HN, and LN, in line with the results from PROGENy where the VEGF pathway was significantly deregulated not only in IgAN, HN and LN, but also in RPGN and DN. 59 different pathways showed significant enrichment in at least 3 CKD entities (Figure 5B). Figure 4B also shows that HN (52), MGN (45), and IgAN (37) are the CKD entities with more pathways differentially enriched in at least 3 entities, a result that agrees with Figure 2B showing these entities in the center of the diffusion map.

### 2.5 Prediction of potential novel drugs that might affect the identified disease signature in different kidney diseases

Finally, we applied a signature-search-engine, L1000CDS^2 24^. L1000CDS^2^ prioritizes small molecules that are expected to have a reverse signature compared to the disease signature. This is based on computing the distance between two signatures of disease data and the LINCS-L1000 data, a large collection of changes in gene expression driven by drugs. We performed this analysis separately for the nine CKD entities and identified 220 small molecules across the CKD entitles (Supplementary Figure 5). In order to narrow down the list of 220 small molecules, we focused on 20 small molecules observed in the L1000CDS^2^ output of at least 3 subtypes (Figure 6A).

**Figure 6.**
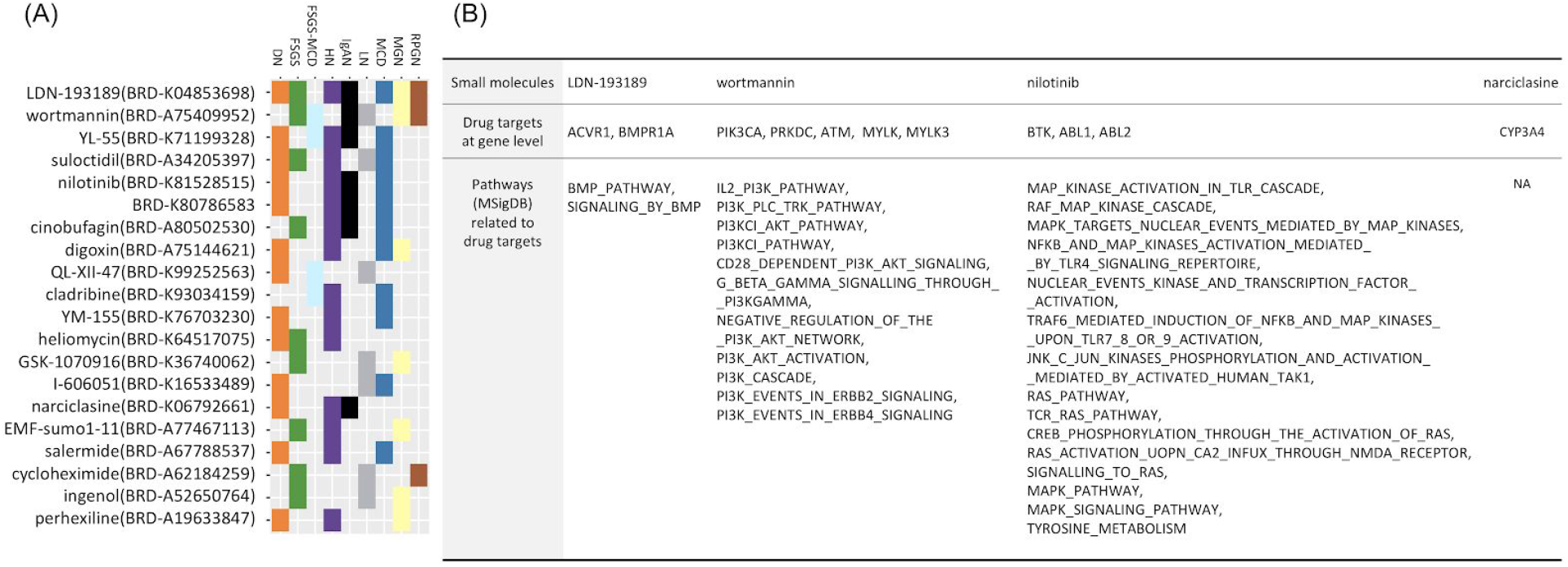
Top 20 Drug Candidates from Drug Repositioning. (A) Distribution of 20 small molecules reversely correlated with at least 3 CKD entities. (B) Table of four small molecules out of the 20 of (A) supported by manual curation. Table shows drugs (first row), protein coding genes targeted by these four drugs (second row) and pathways (MSigDB) related to the biological functions these drugs affect (third row).

By curation of scientific publications, we found that four small molecules have experimental evidence to support their clinical relevance in CKD or renal disease animal model testing (Supplementary Figure 7.). BRD-K04853698 (LDN-193189) which is known as a selective bone morphogenic protein signaling inhibitor, has been shown to suppress endothelial damage in mice with CKD ^25^. Wortmannin, a cell-permeable PI3K inhibitor, decreased albuminuria and podocyte damage in early diabetic nephropathy in rats ^26^. The tyrosine kinase inhibitor nilotinib is used to treat chronic myelogenous leukemia in man.^27^ Nilotinib treatment resulted in stabilized kidney function and prolonged survival after subtotal nephrectomy in rats when compared to vehicle ^28^. Finally, narciclasine was identified and it has been reported to reduce macrophage infiltration and inflammation in the mouse unilateral ureteral obstruction (UUO) model of kidney fibrosis ^29^.

To further explore the association of these drugs with CKD and its’ progression, we analysed the expression data for the targets of the literature supported drug candidates. First, each drug candidate was mapped to genes that encode the proteins targeted by these drugs (Figure 6B). For each gene, its differential expression of any CKD entity against TN was evaluated. Out of the 11 mapped genes, MYLK3, a target of narciclasine, was significantly differentially expressed (under-expressed, logFC<-1, p<0.05) in two CKD entities (IgAN and LN) (Supplementary figure 6). Complementarily, screened drugs were mapped to the pathways they affect based on their functional information. The enrichment of the subset of pathways was evaluated using the previous results from gene set analysis algorithm (piano). This time, only the PI3KCI pathway appeared to be enriched in HN (up-regulated, p<0.05), and as pathway affected by the candidate repositioned drugs (Wortmannin, PI3K inhibitor). Taken together, this data suggests that kidney transcriptomics might be useful to predict potential novel drug candidates.

## 3. Discussion

We have aimed to shed light on the commonalities and differences among glomerular transcriptomes of major kidney diseases contributing to the CKD epidemic affecting >10% of the population worldwide. Multiple pathologies are covered under the broad umbrella of CKD and, while they share a physiological manifestation, i.e. loss of kidney function, the driving molecular process can be different. In this study we explored these processes by analyzing glomerular gene expression data from kidney biopsies obtained via microdissection. We observed expression data of many genes that are considered to be tubule specific in the glomerular dataset e.g. ALB and CALB1. The reason for this might be that microdissection techniques are imperfect, resulting in contamination of glomerular preps with tubule. Current technologies including scRNA-seq will help to dissect expression in particular cell-types of the glomerulus.

Genes such as Quaking (QKI) or Lysozyme C (LYZ), were significantly overexpressed, underexpressed or not altered depending on the underlying kidney disease. It is known that QKI is associated with angiogenic growth factor release and plays a pathological role in the kidney ^30^, while LYZ was known to be related to the extent of vascular damage and heart failure but was recently found to be increased in plasma during CKD progression ^31^. This data supports the fact that despite a stereotypic response of the kidney to injury with glomerulosclerosis, interstitial fibrosis and nephron loss, there are various disease specific differences that are important to understand in order to develop novel personalized therapeutics.

CKD is a complex disease with a high degree of polygenicity. Furthermore, it is a very heterogeneous condition that can be acquired through a variety of biological mechanisms which is reflected by the results of pathway analysis. There was little to no overlap in significantly enriched pathways between the different kidney disease entities. We found 59 different pathways that showed significant enrichment in at least 3 disease entities (Figure 5B), indicating that different disease entities share some general mechanisms but their underlying pathophysiology differs from one entity to another. Besides increasing the interpretability, the pathway analysis identified many more differences among disease-identities than the gene-level analysis (Figure 2A). For example, pathway analysis identified pathways related to the metabolism of lipids and lipoproteins significantly down-regulated in MCD, MGN, and HN; and pathways related to fatty acid metabolism significantly down-regulated in MCD, IgAN, MGN, and HN, results similar to those reported by Kang et al ^6^.

PROGENy (Figure 5A) yielded JAK-STAT, a major cytokine signal transduction regulator ^32^, to be significantly activated in LN with respect to TN and DoROthEA (Figure 3) predicted the TFs IRF1 and STAT1 to be significantly enriched in LN. A pathogenic role of JAK-STAT/STAT1/Interferon signaling in LN is supported by various studies ^33 34 35^.

We also used the signature-matching paradigm to explore potential drugs that could revert the disease phenotype, and found that four drugs hold promise in different CKD entities. Even though more experimental validation is required for the unknown medical interaction between drugs of our results and CKD progression, our approach suggests that it is possible to find promising treatments for CKD via drug repositioning. In particular, for one of the identified drugs, nilotinib, use in humans has already been granted in leukemia and there is supporting data of its value insight at indications for CKD ^28^.

The analysis of the drug targets’ expression found that MYLK3, a gene encoding for one of the targets of narciclasine, was significantly underexpressed in IgAN and LN when compared with TN. Similarly, the PI3KCI pathway, the target of Wortmannin was enriched in HN (up-regulated, p<0.05). This analysis attempted to refine the outcome of the repositioning analysis, and at the same time helped to connect it to the disease mechanism both at the gene as well as the pathway level.

We view our analysis as a first step towards a characterization of the similarities and differences of the various pathologies that lead to CKD. As more data sets become available, either from micro-arrays or RNA-seq, these can be integrated in our pipeline. Furthermore, the burgeoning field of single-cell RNA (scRNA) has just started to produce data sets in kidney ^36,37^ and holds the potential to revolutionize our understanding of the functioning of the kidney and its pathologies ^38 39^. In particular, scRNA data can provide signatures of the many cell types of the kidney, which in turn can be used to deconvolute the composition of cell types^12^ in the more abundant and cost-effective bulk expression datasets ^39^. Other data sets, such as (phospho)proteomics^40^ and metabolomics^41^, may complement gene expression towards a more complete picture of the CKD-entities. Ideally, all these data sets would be collected in a standardized manner to facilitate integration, which was a major hurdle in our study. Such a comprehensive analysis across large cohorts, akin to what has happened for the different tumour types thanks to initiatives such as the International Cancer Genome Consortium, can lead to major improvements in our understanding of and treatment venues for CKD ^42^.

## 4. Methods

### 4.1. Data collection

Raw data CEL files of each microarray dataset - GSE20602 ^10^; GSE32591 ^11^; GSE37460 ^11^; GSE47183 ^12,13^; GSE50469 ^14^ - were downloaded and imported to R (R version 3.3.2). For more information see Supplementary Methods.

### 4.3. Normalization

Cyclic loess normalization was applied using the limma package ^16,43,44^. YuGene transformation was carried out using the YuGene R package ^45^.

### 4.4. Detection of genes with consistently small p-values across all studies

Based on the assumption that common mechanisms might contribute to all CKD entities we performed a Maximum p-value (maxP) method ^46^ - which uses the maximum p-value as the test statistic - on the output of the differential expression analysis of the hypothetically separate studies. For more information see Supplementary Methods.

### 4.6. Diffusion map

The batch mitigated data containing merely the maxP identified (section 4.5.) 1790 genes (Supplementary Material/Data and Code) (FDR < 0.01), were YuGene transformed ^45^ and the destiny R package ^47^ was utilised to produce the diffusion maps.

### 4.7. Functional Analysis

#### 4.7.1. Transcription factor activity analysis

We estimated transcription factor activities in the glomerular CKD entities using DoRothEA^18^ which is a pipeline that tries to estimate transcription factor activity via the expression level of its target genes utilizing a curated database of transcription factor - target gene interactions (TF Regulon). For more information see Supplementary Methods.

#### 4.7.2. Inferring Signaling Pathway Activity fusing PROGENy

We used the cyclic loess normalised and batch effect mitigated expression values for PROGENy ^22^, a method which utilizes downstream gene expression changes due to pathway perturbation in order to infer the upstream signaling pathway activity. For more information see Supplementary Methods.

#### 4.7.3. Pathway Analysis with Piano

Pathway analysis was performed using the piano package from R ^23^. For more information see Supplementary Methods.

### 4.8. Drug repositioning

For each CKD entity, the signature of cosine distances computed by characteristic direction was applied to a signature-search-engine, L1000CDS^2 24^ with the mode of reverse in configuration.

### 4.9. Immunofluorescent staining of human kidney biopsies and analysis

Validation involving human kidney biopsies was approved by the local ethics committee at Karolinska Institutet (Dnr 2017/1991-32). Stainings were performed on 2 μm paraffin-embedded sections as previously described ^48^. For more information see Supplementary Methods.

## Disclosure

The author declare that there is no conflict of interest regarding the publication of this article.

## Acknowledgements

This work was supported by the the JRC for Computational Biomedicine which was partially funded by Bayer AG, the European Union Horizon 2020 grant SyMBioSys MSCA-ITN-2015-ETN #675585 that provided the financial support for A.A and by Grants of the German Research Foundation (KR-4073/3-1, SCHN1188/5-1, SFB/TRR57, SFB/TRR219 TPC01 and C05) a Grant of the European Research Council (ERC-StG 677448), and a Grant of the State of Northrhinewestfalia (Return to NRW) to RK.

## Supplementary Material

**Supplementary Figure 1.**
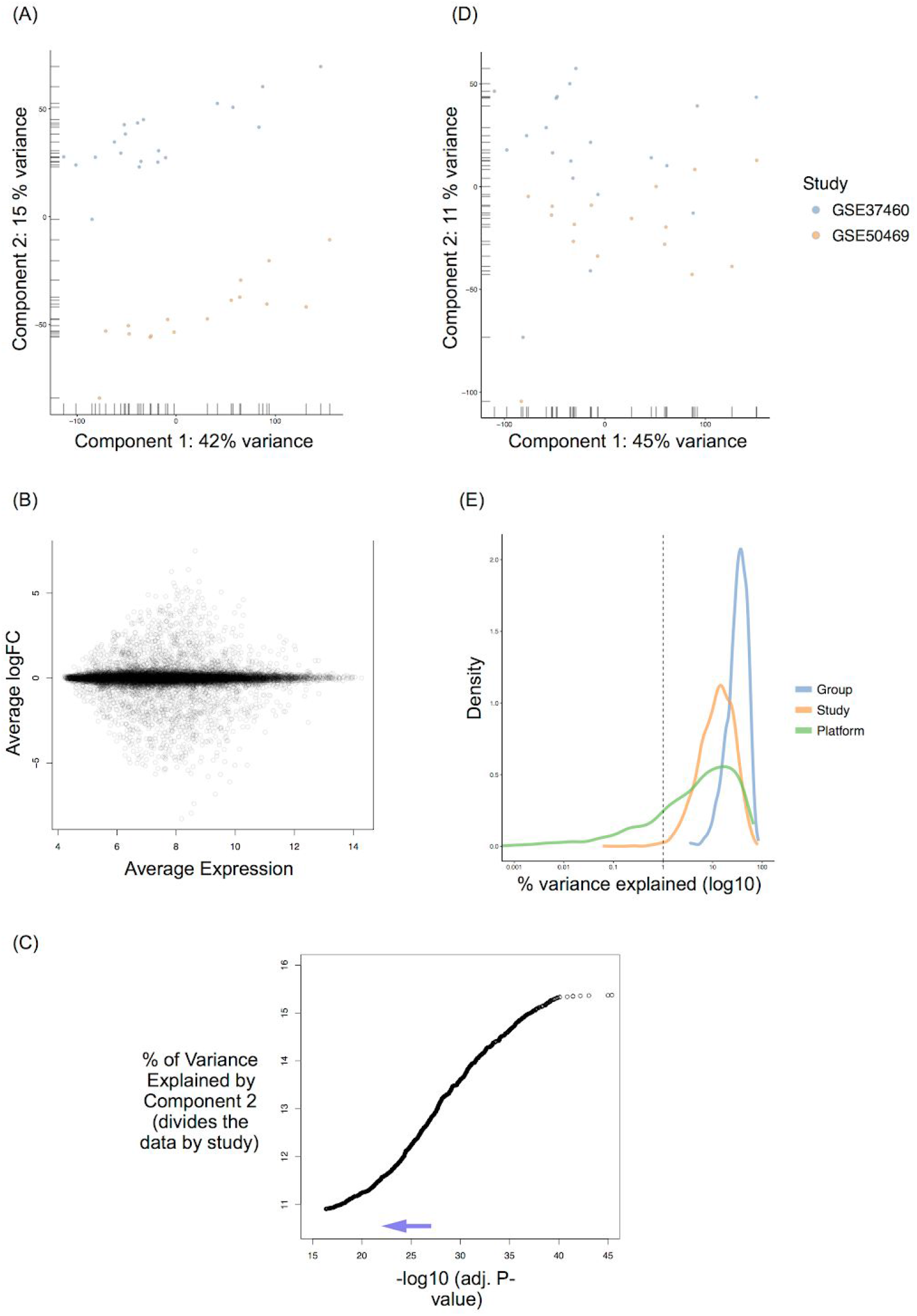
Batch effect mitigation procedure. (A) Principal component analysis (PCA) of gene expression measurements corresponding of IgAN samples from the two distinct studies prior batch effect mitigation. The second principal component separates samples by study. (B) MA plot visualising the difference in gene expression between the GSE37460 and GSE50469 IgAN samples. (C) Percent of variance explained by Principal component 2 (PC2) as a function of the gradual removal of the most affected genes (-log10 adjusted p-value of a particular removed affected gene). (D) PCA of gene expression corresponding to the IgAN samples from the two distinct studies after batch effect mitigation. (E) Depiction of variance for each gene, that is explained by group (CKD entity), study and platform after batch effect mitigation. CKD entity-related variation explains most of the variance in the data.

**Supplementary Figure 2.**
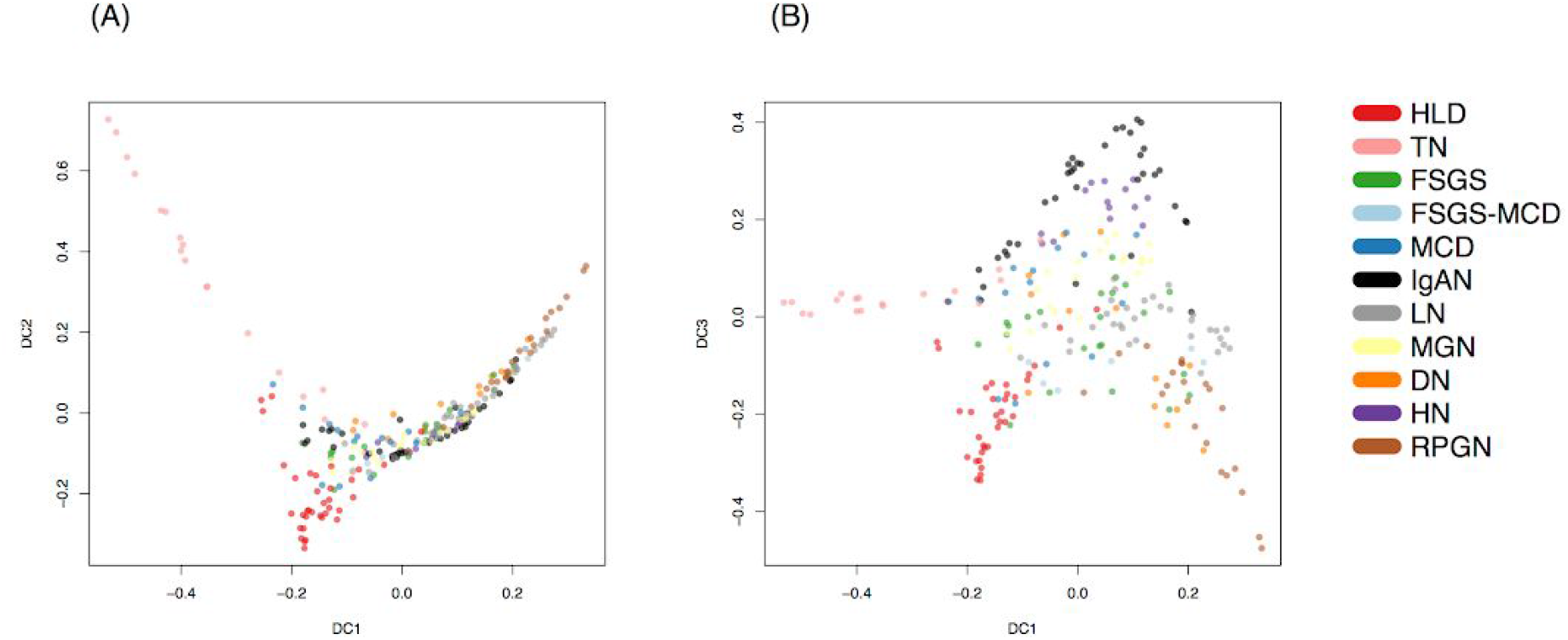
Two dimensional diffusion maps of CKD entities unravel the geometric trajectory of CKD entities based on their comparative transcriptome profile. (A) Dimension component 1 (DC1) is depicted against dimension component 2 (DC2), so that the divergence between the controls and the CKD entities are apparent. (B) DC1 is visualised against dimension component 3 (DC3), revealing the fine distinctions between CKD entities.

**Supplementary Figure 3.**
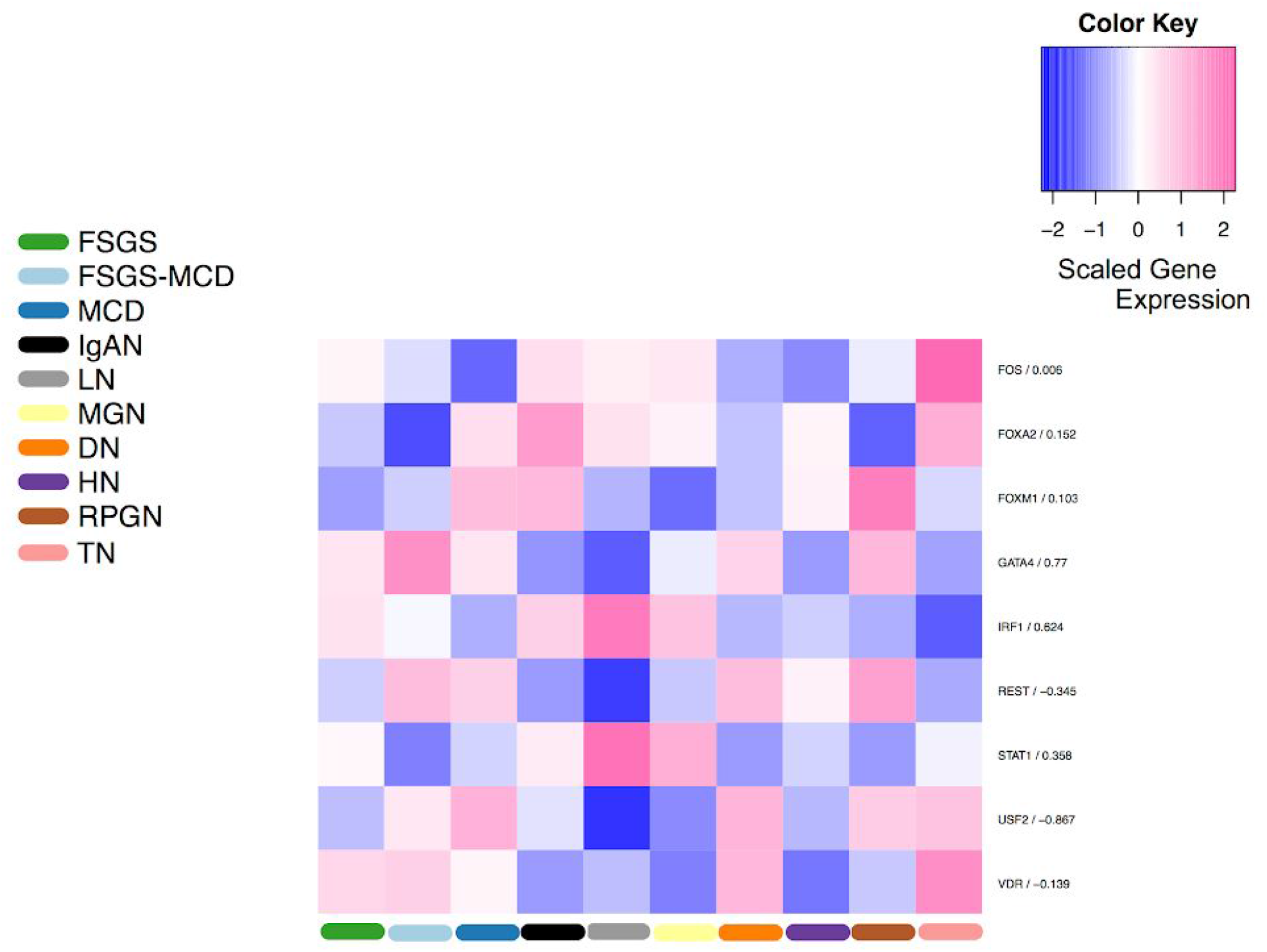
Heatmap depicting the expression of the genes encoding for the transcription factors shown in Figure 3. The expression values were averaged within each condition, then scaled and centered across the conditions. The numbers to the right of factor names are Spearman’s rank-based correlation coefficients of factor activity and factor expression across different CKD entities.

**Supplementary Figure 4.**
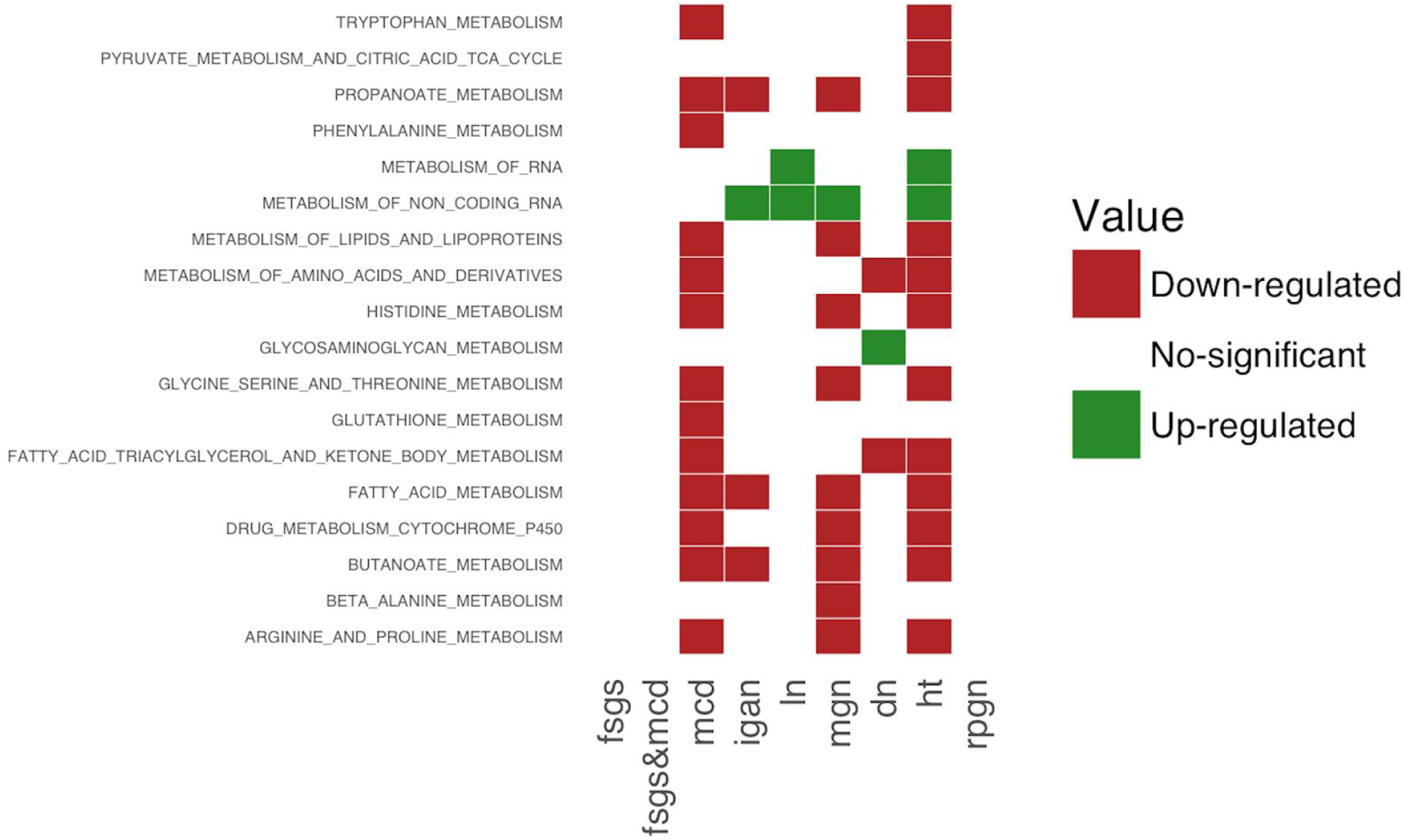
Enrichment of metabolic pathways after gene set analysis. Pathway analysis result in metabolic pathways (‘METABOL’): and their corresponding enrichment: up-regulation (green), down-regulation (red) and non-significant (white). Metabolic pathways are listed in Y axes and disease entities in X axes. Only pathways enriched in at least one disease are shown. Note that FSGS, FSGS-MCD, and RPGN do not have any metabolic pathway significantly affected.

**Supplementary Figure 5.**
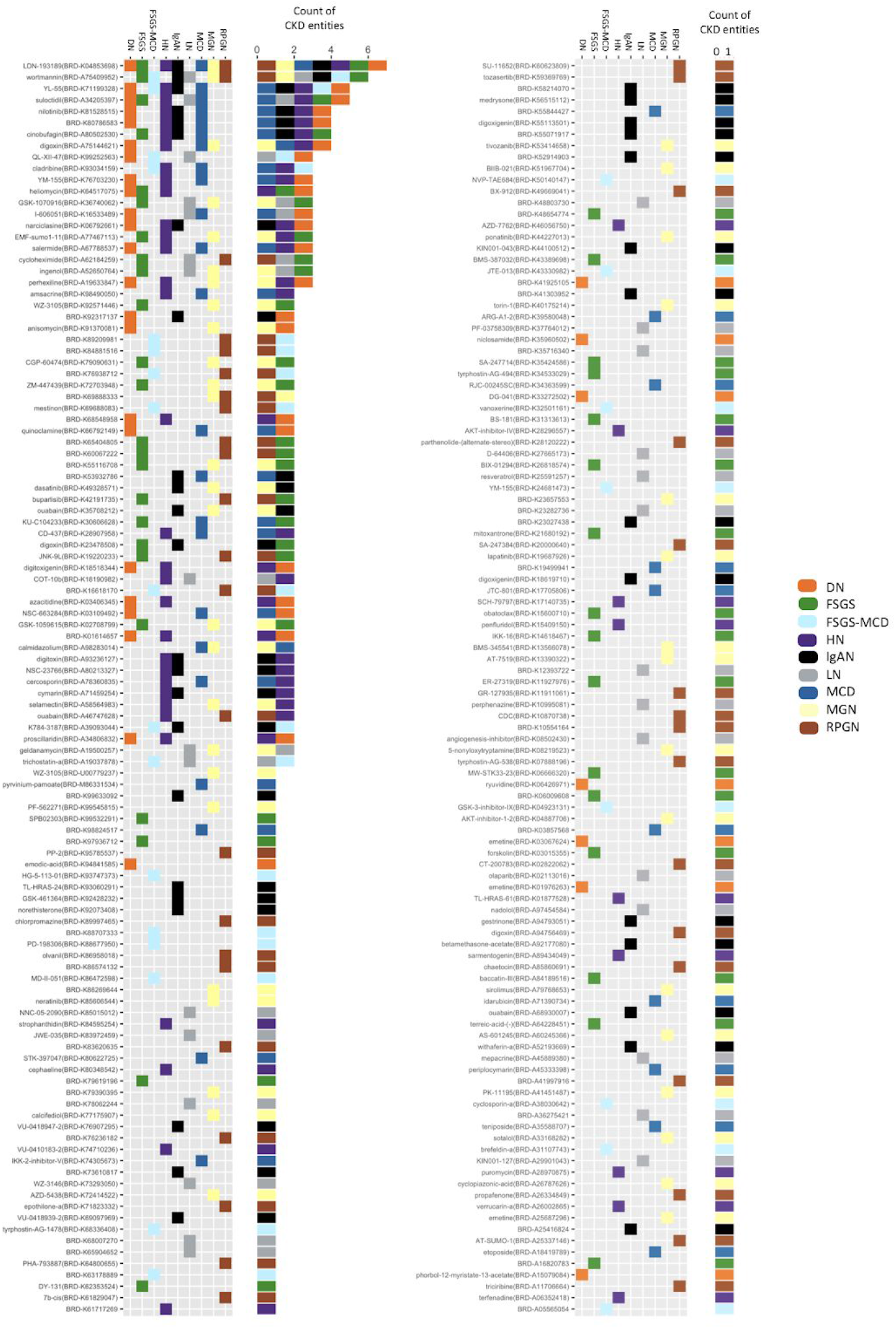
Bar graph (count of CKD entities) and heatmap of the distribution of 220 small molecules reversely correlated with nine CKD entities. Colored bars on both the bar graph and heat map correspond to the subtype of CKD entities and 220 small molecules are represented on the x-axis of both graphs.

**Supplementary Figure 6.**
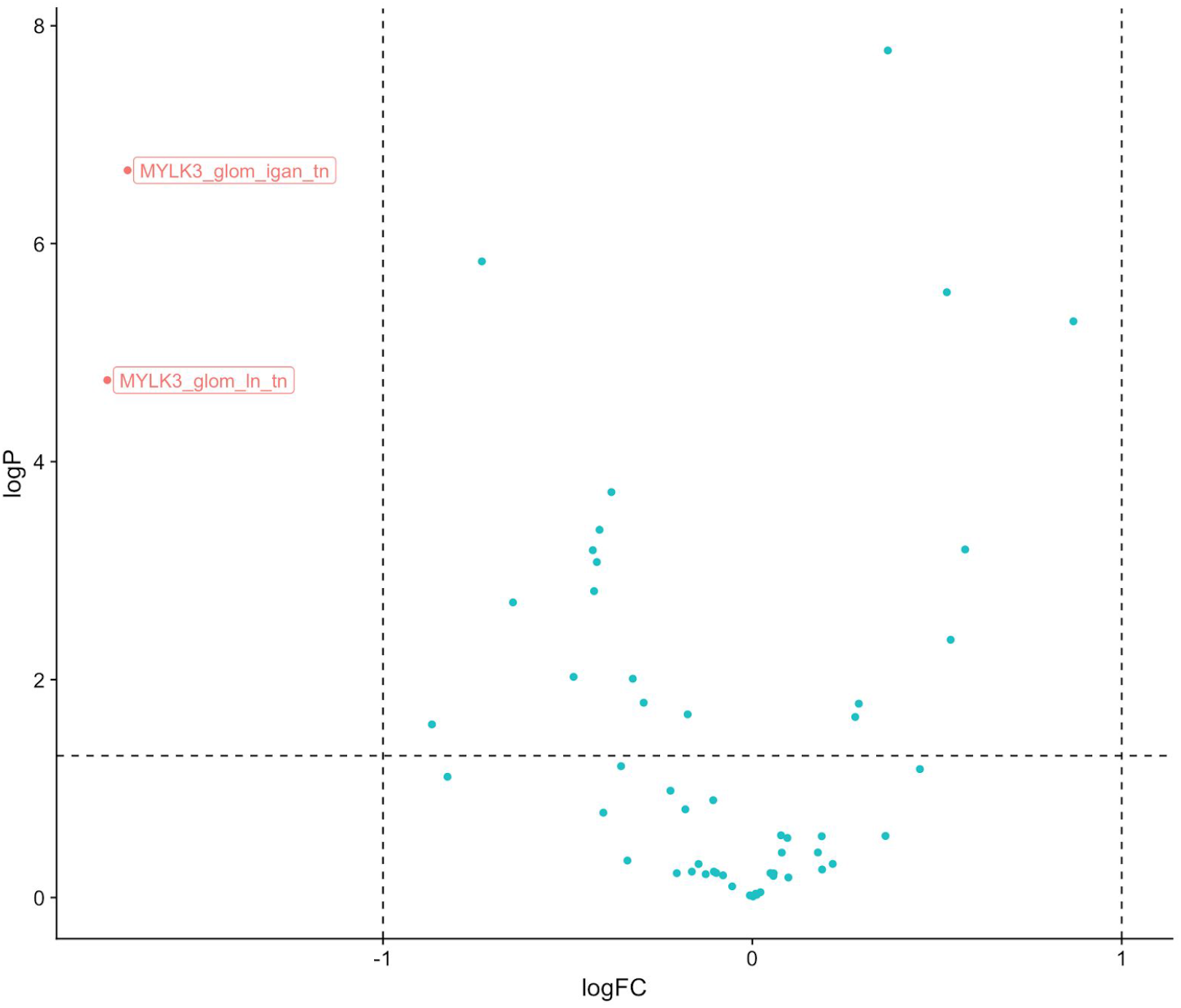
Volcano plot of differential expression of CKD entities vs TN for glomerular samples for the drug targeted genes. X-axis indicates the log2 of the fold change (FC) and the Y-axis the −log10 of the p-value after differential expression analysis using limma.

**Supplementary Figure 7.**
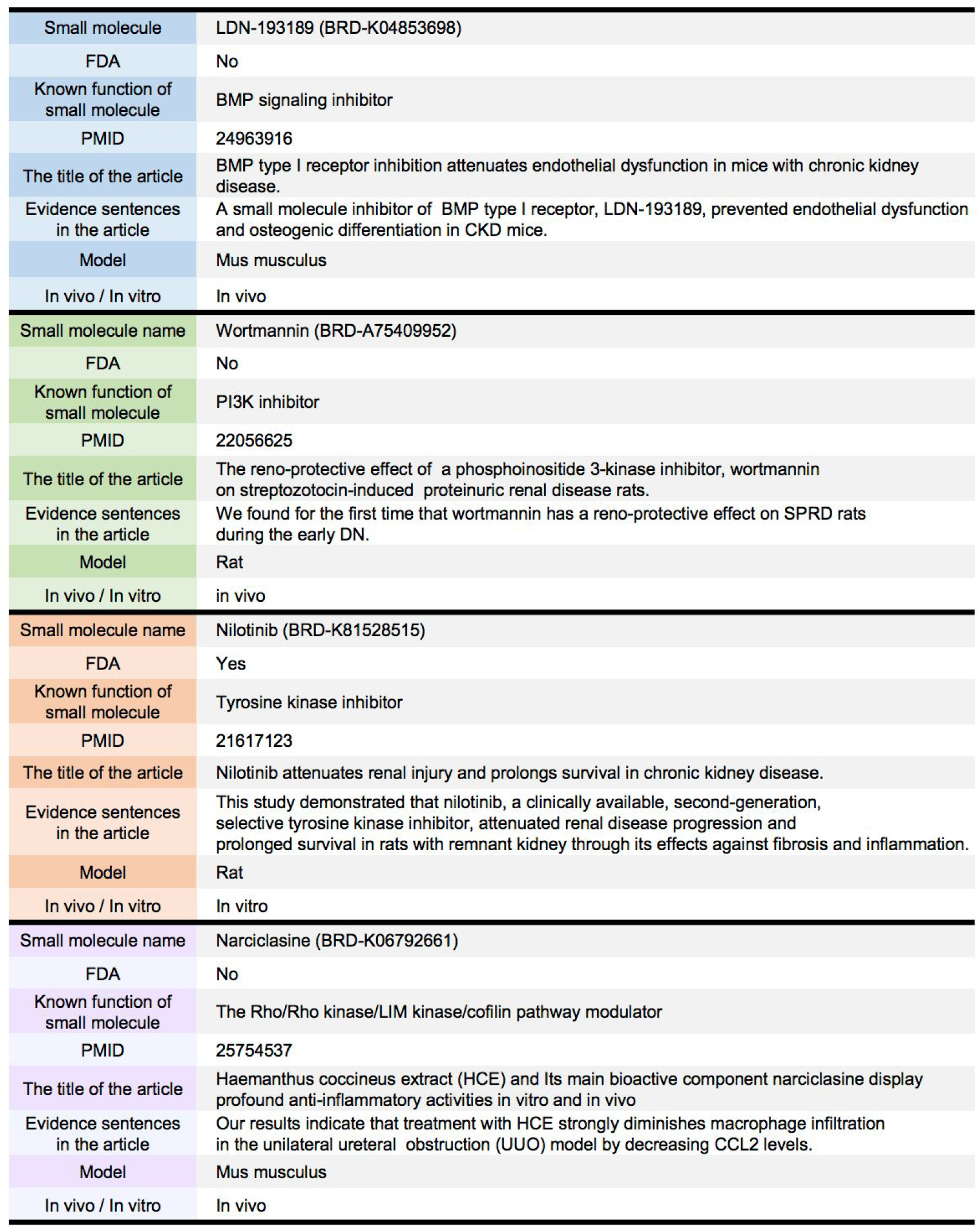
Manual curation of four small molecules. The figure includes drug name corresponding to four small molecules, biological function, FDA approval status and publications describing the clinical relevance of the particular small molecule in CKD.

## Data and Code

https://github.com/saezlab/CKD_Landscape

## Supplementary Methods

### Data collection

Raw data CEL files of each microarray dataset - GSE20602 ^10^; GSE32591 ^11^; GSE37460 ^11^; GSE47183 ^12,13^; GSE50469 ^14^ - were downloaded and imported to R (R version 3.3.2) using the getGEOSuppFiles and read.affy function of the GEOquery and simpleaffy package, respectively ^49^. Each dataset came from either Affymetrix Human Genome U133A Array or Affymetrix Human Genome U133 Plus 2.0 array, therefore the preprocessing was done with the affy R package ^50^ accordingly.

### Preprocessing and mapping

RNA quality was assessed by RNA degradation plots using the AffyRNAdeg function from the affy package. In order to assess the statistical characteristics of the raw data, the affyPLM package ^51^ was used and probe-level metric calculations were carried out on the CEL files by calling the fitPLM function. The homogeneity of probe sets was evaluated by Normalized Unscaled Standard Error (NUSE) and Relative Log Expression (RLE) boxplots ^52–54^. We removed all arrays that showed greater spread of NUSE value distribution with respect to the rest or where the median NUSE value was above 1.05, as these features indicated the sign of low quality array. The RLE values represent the ratio between the expression of a probe set and the median expression of that probe set across all arrays in the data set. The ratios are expected to be centered around zero on a logarithmic scale. RLE boxplots were generated to visualise the distribution of RLE values. Array quality was evaluated by taking NUSE and RLE plots into account. The preprocessing step also constituted background correction and log2 transformation of the raw values, of which was done by the Robust Multichip Average (RMA) package ^55–57^. Probe IDs were mapped to Entrez Gene ID resulting in 20514 (Platform GPL570, Affymetrix Human Genome U133 Plus 2.0 Array) and 12437 (Platform GPL96, Affymetrix Human Genome U133A Array) unique Entrez gene identifiers, respectively. In the case where datasets contained multiple probes for the same Entrez ID gene, the probe with the highest interquartile range (IQR) was retained as the representative of that given gene in the dataset. For this filtering step, the nsFilter function from the genefilter package ^58^ was utilized.

### Correlation of arrays

Correlation of arrays was assessed by hierarchical clustering of the arrays based on gene expression Spearman’s rank-based correlation coefficients. Low Spearman correlation coefficients imply considerable differences between array intensities ^59^.

### Batch effect mitigation

The efficient integration of the data from different sources and platforms requires batch effect management, which should be customised to the data at hand. The current data was heavily affected by platform- and study-specific batch effects, because the outcome categories (CKD entities and their samples) were unevenly distributed across studies and microarray platforms. The commonly used algorithms for correcting batch effects assume a balanced distribution of outcome categories across batches and are vulnerable to the group-batch imbalance ^60–63^. We conducted a stringent batch effect mitigation process in order to minimize the influence of technical heterogeneity. First, we structured the data in a platform-specific manner. Then, we conducted differential gene expression analysis between those identical biological conditions that are originating from distinct study sources after cyclic loess normalization and removed those genes that are significantly differentially expressed between them, as it indicated differences mainly due to the data source, rather than the biological difference. Note that this is a more stringent approach than other batch correction approaches which seek to “model-away” batch-related variance but retain all the data. In our case, this was not possible, and we opted to simply remove genes that are most affected by batch effects. We applied this procedure for the data fragments coming from Affymetrix Human Genome U133 Plus 2.0 Array and Affymetrix Human Genome U133A Array. Next, we merged the data sets between the two platforms using the overlapping genes, followed by a process to mitigate the platform-induced batch effect. This latter procedure is similar to the one used for the data source-specific batch effect mitigation. By applying this stringent procedure, we eliminated the genes that are the most affected by batch effects. For the illustration of this procedure, see Sup. Figure 1. The scater R package ^64^ was used for producing the batch effect management related plots.

### Detection of genes with consistently small p-values across all studies

The maxP test follows a beta distribution that is parametrized by *α* = *K* and *β* = 1 under the null hypothesis:

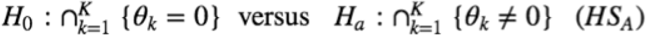

where *θ_k_* is the effect size of study *k*.

This hypothesis setting (symbolised by HS_A_) aims to uncover differentially expressed (DE) genes that acquire non-zero effect sizes in all studies. To phrase it differently, it is designated to unravel DE genes that are characterised by a small p-value across all studies ^46,65,66^.

To obtain the p-values, differential expression analysis was conducted on the batch effect mitigated data using the limma R package ^15^. We contrasted each glomerular CKD entity with tumor nephrectomy condition - each CKD - tumor nephrectomy contrast represented a hypothetically separate “study” - and the lmFit function was used to fit a linear model to the expression data for each probe set in the array series, followed by the estimation of eBayes values and the execution of a moderated t-test by the empirical Bayes method for differential expression (eBayes function) ^15,59^.

### Transcription factor activity analysis with DoRothEA

The cyclic loess normalized expression values of all genes in all conditions were scaled and re-centered across the conditions and the transcription factors activities were estimated from the TF Regulon of DoRothEA^18^ using VIPER ^19^. We then conducted a Spearman’s rank-based correlation between the identified transcription factors’ activity and the scaled and re-centered expression of the genes encoding for these transcription factors. Since as a consequence of the batch effect mitigation procedure we lost many potentially informative genes, the coverage of the TF regulon database was limited and hence our statistical power decrease, meaning that there might be more differentially regulated TFs.

### Inferring Signaling Pathway Activity using PROGENy

The expression values were standardised to express a distance of each CKD sample from tumor nephrectomy (CKD entity sample scaled by the standard deviation of tumor nephrectomy). We used the overlapping genes between the standardised gene expression matrix and the PROGENy model. Then, a matrix multiplication was done to get the product matrix, containing a PROGENy score for each pathway in each CKD entity. A positive PROGENy score in a given pathway in a given CKD entity implies higher signaling activity compared to that specific pathways’ activity in tumor nephrectomy, and vice versa for a negative PROGENy score.

Statistical significance was assessed using permutation-based hypothesis testing. We resampled the standardised gene expression values in a way that results in the randomised allocation of expression values to different glomerular disease labels. This resampling was done ten thousand times. We then computed PROGENy scores from these permuted Z-scores, resulting in a list of glomerular CKD entity specific PROGENy scores. By applying this approach we generated an empirical null distribution on the basis of the original gene expression sample distribution. The probability that the original PROGENy score in a given glomerular CKD entity is coming from the estimated null distribution or not was evaluated in a pathway-specific manner. We used a p-value of 0.05 as the threshold for statistical significance. Furthermore, we applied the Benjamini-Hochberg adjustment ^67^ on the p-values to correct for multiple testing.

### Pathway Analysis with Piano

Pathway analysis was performed using the piano package from R ^23^. The Molecular Signature Database - Curated Pathways - Canonical Pathways (MSigDB-C2-CP) was used as biological model to map the individual genes to functional sets. Gene-level statistics were obtained after applying the limma algorithm (see section 4.5.). All disease entities were compared to tumor nephrectomy, because the healthy living donor samples were highly corrupted by batch effects and as a result of the batch effect mitigation, we had to remove a considerably large number of genes from these samples. The following ten methods (with their corresponding gene-level statistics) were used as input of the pathway analysis algorithm to calculate gene set enrichment: Fisher (PVal), Stouffer (PVal), Reporter (PVal), PAGE (TVal), Tail Strength (PVal), GSEA (TVal), Mean (FC), Median (FC), Sum (FC), MaxMean (TVal). For each pathway/p-value pair the geometrical average across all ten methods was calculated. For simplicity, only the p_dist_down and p_dist_up features were retrieved.

### Drug repositioning

Cyclic loess normalized gene expression data for nine glomerular CKD entities were analyzed separately for measuring characteristic direction (CD) ^68^. Cosine distance for each gene was computed to the line which has 90 degree to the hyperplane which set the given CKD entity apart from tumor nephrectomy in N-dimensional gene expression space. Then, for each CKD entity, the signature of cosine distances computed by characteristic direction was applied to a signature-search-engine, L1000CDS^2 24^ with the mode of reverse in configuration. L1000CDS^2^ provided the top 50 ranked small molecule candidates with 1-cos(a), p-value, drug database links. Significant small molecules with FDR < 0.05 were filtered in for the nine CKD entities, separately. For converting the name of small molecules into general chemical names, we referred to LINCS phase I, II dataset stored in GEO (GSE92742, GSE70138) ^69^.

### Immunofluorescent staining of human kidney biopsies and analysis

The following primary antibodies were used and incubated 1h: USF-2 (5E9, Santa Cruz, 1:100, mouse), FOXM1 (ab207298, Abcam, 1:100, rabbit), CD44-Alexa-Fluor-647 (IM7, BioLegend, 1:100, rat). The following secondary antibodies were used: donkey anti**–**rabbit, **–**mouse, Alexa Fluor 546 (Dianova, 1:200). The nuclei were stained using DAPI (Sigma). Sections were evaluated with a Nikon A1R confocal microscope with 40x objective. Image processing was performed using NIH ImageJ software.

## References

1. Hill NR, Fatoba ST, Oke JL et al. Global Prevalence of Chronic Kidney Disease - A Systematic Review and Meta-Analysis. PLoS One 2016; 11: e0158765.

2. Hamer RA, El Nahas AM. The burden of chronic kidney disease. BMJ 2006; 332: 563–564.

3. Beckerman P, Qiu C, Park J et al. Human Kidney Tubule-Specific Gene Expression Based Dissection of Chronic Kidney Disease Traits. EBioMedicine 2017; 24: 267–276.

4. Nair V, Komorowsky CV, Weil EJ et al. A molecular morphometric approach to diabetic kidney disease can link structure to function and outcome. Kidney Int. 2017.

5. Schena FP, Nistor I, Curci C. Transcriptomics in kidney biopsy is an untapped resource for precision therapy in nephrology: a systematic review. Nephrol. Dial. Transplant 2017.

6. Kang HM, Ahn SH, Choi P et al. Defective fatty acid oxidation in renal tubular epithelial cells has a key role in kidney fibrosis development. Nat. Med. 2015; 21: 37–46.

7. Ju W, Nair V, Smith S et al. Tissue transcriptome-driven identification of epidermal growth factor as a chronic kidney disease biomarker. Sci. Transl. Med. 2015; 7: 316ra193.

8. Edgar R, Domrachev M, Lash AE. Gene Expression Omnibus: NCBI gene expression and hybridization array data repository. Nucleic Acids Res. 2002; 30: 207–210.

9. Barrett T, Wilhite SE, Ledoux P et al. NCBI GEO: archive for functional genomics data sets--update. Nucleic Acids Res. 2013; 41: D991–5.

10. Neusser MA, Lindenmeyer MT, Moll AG et al. Human nephrosclerosis triggers a hypoxia-related glomerulopathy. Am. J. Pathol. 2010; 176: 594–607.

11. Berthier CC, Bethunaickan R, Gonzalez-Rivera T et al. Cross-species transcriptional network analysis defines shared inflammatory responses in murine and human lupus nephritis. J. Immunol. 2012; 189: 988–1001.

12. Ju W, Greene CS, Eichinger F et al. Defining cell-type specificity at the transcriptional level in human disease. Genome Res. 2013; 23: 1862–1873.

13. Martini S, Nair V, Keller BJ et al. Integrative biology identifies shared transcriptional networks in CKD. J. Am. Soc. Nephrol. 2014; 25: 2559–2572.

14. Hodgin JB, Berthier CC, John R et al. The molecular phenotype of endocapillary proliferation: novel therapeutic targets for IgA nephropathy. PLoS One 2014; 9: e103413.

15. Phipson B, Lee S, Majewski IJ et al. ROBUST HYPERPARAMETER ESTIMATION PROTECTS AGAINST HYPERVARIABLE GENES AND IMPROVES POWER TO DETECT DIFFERENTIAL EXPRESSION. Ann. Appl. Stat. 2016; 10: 946–963.

16. Ritchie ME, Phipson B, Wu D et al. limma powers differential expression analyses for RNA-sequencing and microarray studies. Nucleic Acids Res. 2015; 43: e47.

17. Haghverdi L, Buettner F, Theis FJ. Diffusion maps for high-dimensional single-cell analysis of differentiation data. Bioinformatics 2015; 31: 2989–2998.

18. Garcia-Alonso LM, Iorio F, Matchan A et al. Transcription factor activities enhance markers of drug sensitivity in cancer. Cancer Res. 2017.

19. Alvarez MJ, Shen Y, Giorgi FM et al. Functional characterization of somatic mutations in cancer using network-based inference of protein activity. Nat. Genet. 2016; 48: 838–847.

20. Matsuda M, Tamura K, Wakui H et al. Upstream stimulatory factors 1 and 2 mediate the transcription of angiotensin II binding and inhibitory protein. J. Biol. Chem. 2013; 288: 19238–19249.

21. Gentles AJ, Newman AM, Liu CL et al. The prognostic landscape of genes and infiltrating immune cells across human cancers. Nat. Med. 2015; 21: 938–945.

22. Schubert M, Klinger B, Klünemann M et al. Perturbation-response genes reveal signaling footprints in cancer gene expression. Nat. Commun. 2018; 9: 20.

23. Väremo L, Nielsen J, Nookaew I. Enriching the gene set analysis of genome-wide data by incorporating directionality of gene expression and combining statistical hypotheses and methods. Nucleic Acids Res. 2013; 41: 4378–4391.

24. Duan Q, Reid SP, Clark NR, et al. L1000CDS2: LINCS L1000 characteristic direction signatures search engine. NPJ Syst Biol Appl 2016; 2.

25. Kajimoto H, Kai H, Aoki H et al. BMP type I receptor inhibition attenuates endothelial dysfunction in mice with chronic kidney disease. Kidney Int. 2015; 87: 128–136.

26. Kim SH, Jang YW, Hwang P et al. The reno-protective effect of a phosphoinositide 3-kinase inhibitor wortmannin on streptozotocin-induced proteinuric renal disease rats. Exp. Mol. Med. 2012; 44: 45–51.

27. Blay J-Y, von Mehren M. Nilotinib: a novel, selective tyrosine kinase inhibitor. Semin. Oncol. 2011; 38 Suppl 1: S3–9.

28. Iyoda M, Shibata T, Hirai Y et al. Nilotinib attenuates renal injury and prolongs survival in chronic kidney disease. J. Am. Soc. Nephrol. 2011; 22: 1486–1496.

29. Fuchs S, Hsieh LT, Saarberg W et al. Haemanthus coccineusextract and its main bioactive component narciclasine display profound anti-inflammatory activitiesin vitroandin vivo. J. Cell. Mol. Med. 2015; 19: 1021–1032.

30. Gomez IG, Nakagawa N, Duffield JS. MicroRNAs as novel therapeutic targets to treat kidney injury and fibrosis. Am. J. Physiol. Renal Physiol. 2016; 310: F931–44.

31. Glorieux G, Mullen W, Duranton F et al. New insights in molecular mechanisms involved in chronic kidney disease using high-resolution plasma proteome analysis. Nephrol. Dial. Transplant 2015; 30: 1842–1852.

32. Rawlings JS, Rosler KM, Harrison DA. The JAK/STAT signaling pathway. J. Cell Sci. 2004; 117: 1281–1283.

33. Dong J, Wang QX, Zhou CY et al. Activation of the STAT1 signalling pathway in lupus nephritis in MRL/lpr mice. Lupus 2007; 16: 101–109.

34. Wang S, Yang N, Zhang L et al. Jak/STAT signaling is involved in the inflammatory infiltration of the kidneys in MRL/lpr mice. Lupus 2010; 19: 1171–1180.

35. Ripoll È, de Ramon L, Draibe Bordignon J et al. JAK3-STAT pathway blocking benefits in experimental lupus nephritis. Arthritis Res. Ther. 2016; 18: 134.

36. Sivakamasundari V, Bolisetty M, Sivajothi S. Comprehensive Cell Type Specific Transcriptomics of the Human Kidney. bioRxiv 2017.

37. Wu H, Uchimura K, Donnelly E et al. Comparative analysis of kidney organoid and adult human kidney single cell and single nucleus transcriptomes. bioRxiv 2017.

38. Wu H, Humphreys BD. The promise of single-cell RNA sequencing for kidney disease investigation. Kidney Int. 2017; 92: 1334–1342.

39. Kiryluk K, Bomback AS, Cheng Y-L et al. Precision Medicine for Acute Kidney Injury (AKI): Redefining AKI by Agnostic Kidney Tissue Interrogation and Genetics. Semin. Nephrol. 2018; 38: 40–51.

40. Liu P, Lassén E, Nair V et al. Transcriptomic and Proteomic Profiling Provides Insight into Mesangial Cell Function in IgA Nephropathy. J. Am. Soc. Nephrol. 2017; 28: 2961–2972.

41. Hocher B, Adamski J. Metabolomics for clinical use and research in chronic kidney disease. Nat. Rev. Nephrol. 2017; 13: 269–284.

42. Mariani LH, Pendergraft WF 3rd, Kretzler M. Defining Glomerular Disease in Mechanistic Terms: Implementing an Integrative Biology Approach in Nephrology. Clin. J. Am. Soc. Nephrol. 2016; 11: 2054–2060.

43. Smyth GK. limma: Linear Models for Microarray Data. In: Bioinformatics and Computational Biology Solutions Using R and Bioconductor. Springer, New York, NY, 2005, pp.397–420.

44. Smyth GK, Speed T. Normalization of cDNA microarray data. Methods 2003; 31: 265–273.

45. Lê Cao K-A, Rohart F, McHugh L et al. YuGene: a simple approach to scale gene expression data derived from different platforms for integrated analyses. Genomics 2014; 103: 239–251.

46. Wilkinson B. A statistical consideration in psychological research. Psychol. Bull. 1951; 48: 156–158.

47. Angerer P, Haghverdi L, Büttner M et al. destiny: diffusion maps for large-scale single-cell data in R. Bioinformatics 2016; 32: 1241–1243.

48. Kuppe C, Gröne H-J, Ostendorf T et al. Common histological patterns in glomerular epithelial cells in secondary focal segmental glomerulosclerosis. Kidney Int. 2015; 88: 990–998.

49. Wilson CL, Miller CJ. Simpleaffy: a BioConductor package for Affymetrix Quality Control and data analysis. Bioinformatics 2005; 21: 3683–3685.

50. Gautier L, Cope L, Bolstad BM et al. affy--analysis of Affymetrix GeneChip data at the probe level. Bioinformatics 2004; 20: 307–315.

51. Bolstad BM. affyPLM: Methods for fitting probe-level models. BioConductor version 2. 0 package 2007.

52. Bolstad BM. Low-level Analysis of High-density Oligonucleotide Array Data: Background, Normalization and Summarization. 2004. University of California, Berkeley.(http://bmbolstad.com.

53. Bolstad BM, Collin F, Brettschneider J et al. Quality Assessment of Affymetrix GeneChip Data. In: Bioinformatics and Computational Biology Solutions Using R and Bioconductor. Springer, New York, NY, 2005, pp.33–47.

54. Brettschneider J, Collin F, Bolstad BM et al. Quality Assessment for Short Oligonucleotide Microarray Data. Technometrics 2008; 50: 241–264.

55. Irizarry RA, Bolstad BM, Collin F et al. Summaries of Affymetrix GeneChip probe level data. Nucleic Acids Res. 2003; 31: e15.

56. Irizarry RA, Hobbs B, Collin F et al. Exploration, normalization, and summaries of high density oligonucleotide array probe level data. Biostatistics 2003; 4: 249–264.

57. Bolstad BM, Irizarry RA, Astrand M et al. A comparison of normalization methods for high density oligonucleotide array data based on variance and bias. Bioinformatics 2003; 19: 185–193.

58. Gentleman R, Carey V, Huber W et al. Genefilter: Methods for filtering genes from microarray experiments. R package version 2011.

59. Baetke SC, Adriaens ME, Seigneuric R et al. Molecular pathways involved in prostate carcinogenesis: insights from public microarray datasets. PLoS One 2012; 7: e49831.

60. Johnson WE, Li C, Rabinovic A. Adjusting batch effects in microarray expression data using empirical Bayes methods. Biostatistics 2007; 8: 118–127.

61. Goh WWB, Wang W, Wong L. Why Batch Effects Matter in Omics Data, and How to Avoid Them. Trends Biotechnol. 2017; 35: 498–507.

62. Nygaard V, Rødland EA, Hovig E. Methods that remove batch effects while retaining group differences may lead to exaggerated confidence in downstream analyses. Biostatistics 2016; 17: 29–39.

63. Leek JT, Scharpf RB, Bravo HC et al. Tackling the widespread and critical impact of batch effects in high-throughput data. Nat. Rev. Genet. 2010; 11: 733.

64. McCarthy DJ, Campbell KR, Lun ATL et al. Scater: pre-processing, quality control, normalization and visualization of single-cell RNA-seq data in R. Bioinformatics 2017; 33: 1179–1186.

65. Chang L-C, Lin H-M, Sibille E et al. Meta-analysis methods for combining multiple expression profiles: comparisons, statistical characterization and an application guideline. BMC Bioinformatics 2013; 14: 368.

66. Song C, Tseng GC. HYPOTHESIS SETTING AND ORDER STATISTIC FOR ROBUST GENOMIC META-ANALYSIS. Ann. Appl. Stat. 2014; 8: 777–800.

67. Benjamini Y, Hochberg Y. Controlling the False Discovery Rate: A Practical and Powerful Approach to Multiple Testing. J. R. Stat. Soc. Series B Stat. Methodol. 1995; 57: 289–300.

68. Clark NR, Hu KS, Feldmann AS et al. The characteristic direction: a geometrical approach to identify differentially expressed genes. BMC Bioinformatics 2014; 15: 79.

69. Subramanian A, Narayan R, Corsello SM et al. A Next Generation Connectivity Map: L1000 Platform and the First 1,000,000 Profiles. Cell 2017; 171: 1437–1452.e17.

